# Regulatory architecture of the RCA gene cluster captures an intragenic TAD boundary, CTCF-mediated chromatin looping and a long-range intergenic enhancer

**DOI:** 10.1101/2020.02.16.941070

**Authors:** Jessica Cheng, Joshua S. Clayton, Rafael D. Acemel, Ye Zheng, Rhonda L. Taylor, Sündüz Keleş, Martin Franke, Susan A. Boackle, John B. Harley, Elizabeth Quail, José L. Gómez-Skarmeta, Daniela Ulgiati

## Abstract

The Regulators of Complement Activation (RCA) gene cluster comprises several tandemly arranged genes with shared functions within the immune system. RCA members, such as complement receptor 2 (*CR2*), are well-established susceptibility genes in complex autoimmune diseases. Altered expression of RCA genes has been demonstrated at both the functional and genetic level, but the mechanisms underlying their regulation are not fully characterised. We aimed to investigate the structural organisation of the RCA gene cluster to identify key regulatory elements that influence the expression of *CR2* and other genes in this immunomodulatory region. Using 4C, we captured extensive CTCF-mediated chromatin looping across the RCA gene cluster in B cells and showed these were organised into two topologically associated domains (TADs). Interestingly, an inter-TAD boundary was located within the *CR1* gene at a well-characterised segmental duplication. Additionally, we mapped numerous gene-gene and gene-enhancer interactions across the region, revealing extensive co-regulation. Importantly, we identified an intergenic enhancer and functionally demonstrated this element upregulates two RCA members (*CR2* and *CD55*) in B cells. We have uncovered novel, long-range mechanisms whereby autoimmune disease susceptibility may be influenced by genetic variants, thus highlighting the important contribution of chromatin topology to gene regulation and complex genetic disease.

**SIGNIFICANCE:** The complement system is a complex network of protein effectors and regulators that play a key role in immunity. Several regulators of complement response are clustered within Regulators of Complement Activation (RCA) gene family. Its members are all functionally, structurally, and genetically related. However, the functional relevance of this close gene organisation is unknown. We show that the clustering of the RCA members is due to shared long-range regulatory elements and physical chromatin looping. We also reveal that the RCA genes are divided into two adjacent chromatin domains and a domain boundary falls within the body of an expressed gene (*CR1*). Overall, our findings in the RCA cluster offer insights into their evolution, biology and roles in disease.

## INTRODUCTION

The complement system is a major immune network of soluble proteins and membrane receptors which elicit potent, innate responses against pathogens, immune complexes and apoptotic cells (1). The complement system is activated by one of three major pathways (classical, alternative or lectin), triggering a series of proteolytic cleavage events which ultimately converge to form the C3 convertase. The C3 convertase enzyme catalyses, in part, the formation of complement effector peptides (C3a, C5a, C3b and C5b) which mediate local inflammation, cell lysis and cell clearance (1). Additionally, complement components are capable of binding numerous immune cell types and activating other immune pathways, including adaptive B cell and T cell responses (2, 3). Complement therefore represents an important bridge between the innate and adaptive immune systems and allows for effective co-ordination of immune responses (4).

The complement cascade is intricately controlled to ensure a sufficient immune response is generated while preventing damage to self (1). In humans, a number of these regulatory proteins are located in a gene cluster known as the Regulators of Complement Activation (RCA) on chromosome 1q32.2. This includes the plasma protein C4 binding protein (encoded by alpha (*C4BPA*) and beta (*C4BPB*) subunits), and several membrane receptors; decay-accelerating factor (DAF, *CD55*), complement receptors 2 and 1 (*CR2* and *CR1*), and membrane co-factor protein (MCP, *CD46*) (5). Several duplicated pseudogenes within the RCA cluster have also been identified (6, 7) of which CR1-like (*CR1L)* has been best characterised (8). All members of the RCA gene cluster are composed of tandem 60 – 70 amino acid motifs known as short consensus repeats (SCRs) which bind complement components and primarily regulate the complement response through inhibition or activation of C3 convertase (1, 5). As such, this gene cluster is believed to have been derived from complex duplications of a common ancestral gene, followed by the diversification of function (5). In addition to their important roles in innate immune responses, members of RCA gene cluster are involved in the processes of tissue injury, inflammation and apoptosis. Accordingly, they have been implicated in a range of inflammatory and autoimmune disorders (9-12).

A role for complement receptors CR2 and CR1 in the autoimmune disease, Systemic Lupus Erythematosus (SLE) is well established. SLE is characterised by the presence of antibodies directed against nuclear antigens and has a complex aetiology with a strong genetic component (13, 14). CR2 and CR1 regulate B cell responses by modulating B cell activation and antibody production upon binding of complement-tagged antigens (15, 16). Aberrant expression of CR2 on the surface of B cells has been demonstrated both in mouse models of the disease (17, 18) and SLE patients (19, 20), which functionally contributes to B cell autoreactivity and autoimmune disease susceptibility (21-23). The *CR1* gene contains an 18 kb intragenic segmental duplication, known as ‘low copy repeat 1’ (LCR1). This repeat results in different structural alleles of *CR1* with recognised association to SLE susceptibility, but the functional role of this large genomic duplication is not well understood (24). The *CR2* gene has also been implicated in SLE at the genetic level through linkage analyses (25-27) and association studies (27-29). A SNP within the first intron of *CR2* (rs1876453) was shown to alter the expression of the neighbouring gene (*CR1*) without influencing CR2 expression (29), indicating that expression of these genes in the RCA cluster may be co-regulated. Functionally, rs1876453 was shown to influence the binding affinity of CCCTC-binding factor (CTCF) to *CR2*, suggesting that CTCF may have a role in co-regulating expression of *CR2, CR1* and the RCA gene cluster (29).

CTCF is an important transcription factor which was first identified as an insulator of gene expression, and is now known to have several roles in gene regulation. Additionally, CTCF has been shown to play a critical role in forming chromatin loops and mediating interactions between distal loci (30). Chromatin loops are organised into genomic compartments known as topologically associated domains (TADs) (31). The current model proposed to explain TAD formation involves CTCF and the cohesin complex, whereby loops are dynamically formed through ‘loop extrusion’ between distal CTCF sites in convergent or ‘forward-facing’ orientation (32). TADs are recognised to be constitutively maintained in different cell types but may alternate between active (“A”) and inactive (“B”) compartment types depending on the cellular context (33, 34). While genes within the same TAD tend to be co-expressed, not all genes within a TAD are necessarily expressed simultaneously. Rather, in a given context, TADs restrict chromatin interactions between genes and distal regulatory elements, such as enhancers, to ensure that gene expression is properly controlled (33, 35).

Enhancers represent an important class of distal-regulatory elements which are largely responsible for governing cell-type specific gene expression patterns. Enhancers bind transcription factors to upregulate expression of genes and are located distal to gene promoters in the linear genome but are positioned in close proximity by chromatin looping (36). Importantly, the majority of disease-associated SNPs from genome-wide association studies (GWAS) fall within enhancer regions (37). Enhancer elements have been predicted in the genome by the presence of epigenetic marks such as enrichment of H3K27ac and expression of short, bi-directional transcripts termed enhancer RNA (eRNA) (38, 39). However, enhancers can simultaneously regulate expression of multiple genes, regulate genes large distances away and skip their neighbouring gene/s, which has hindered the identification of their target gene/s (36). The mapping of chromatin interactions through high-throughput chromatin conformation capture technologies, such as Hi-C and capture Hi-C (CHi-C), has aided in the identification of enhancer targets. However, these data are still limited by resolution and the physical chromatin interactions detected using these methods may not necessarily be functional. As such, experimental validation of enhancers and physically associating enhancer-gene pairs is imperative to determine their influence on gene expression (36, 40). In addition, large repetitive regions in the genome, such as the LCR in *CR1*, cannot be uniquely aligned and readily analysed using next-generation sequencing technologies. As a result, these regions are under-represented in high-throughput epigenetic and chromatin conformation capture datasets.

The aim of this investigation was to explore the structural organisation of the RCA gene cluster in order to identity transcriptional elements which may co-regulate the expression of genes in this important immunomodulatory cluster. In this study, we examined genomic interactions across the RCA gene cluster using chromosome conformation capture and showed that long-range chromatin interactions are involved in the co-regulation and co-expression of several RCA members in the B cell lineage. Further, we identified an intragenic TAD boundary which discretely separates chromatin interactions in the RCA gene cluster into two domains and co-localises to the intragenic segmental duplication in *CR1*. Importantly, we functionally interrogated a putative long-range enhancer and demonstrated that it co-regulates two genes within a TAD in B cells. Collectively, we have revealed how three-dimensional chromatin organisation plays an important role in regulating the RCA gene cluster and have uncovered novel regulatory loci which govern the expression of these genes.

## RESULTS

### Chromatin interactions within the RCA gene cluster are organised into two TADs

To investigate the structural arrangement of the RCA gene cluster in B cells, we examined raw Hi-C data in the GM12878 B lymphoblastoid cell line from Rao *et al*. (41) at 10 kb resolution. The intergenic region between *CD55* and *CR2*, and loci across the complement receptor genes (*CR2* and *CR1*) engaged in highly frequent interactions with loci more than 400 kb upstream near *C4BPB* and *C4BPA* (**Figure 1A**). No notable interactions between these regions with downstream genes *CR1L* and *CD46* were observed (**Figure 1A**), indicating that chromatin interactions in this region may be directionally constrained and organised to more than one TAD. This pattern of interaction was consistent across the 6 other cell lines also examined at 10 kb resolution in Rao *et al*. (41) Hi-C dataset (K562, HMEC, NHEK, IMR90, KBM7, HUVEC) (**Supplementary Figure 1**). The *CR1* intragenic duplication (*CR1* exon 5 – 20) is included on the human reference genome, but Hi-C interaction data was filtered out at this region in all datasets due to sequence repetitiveness and sequence unmappability (**Figure 1A, Supplementary Figure 1**).

**Figure 1:**
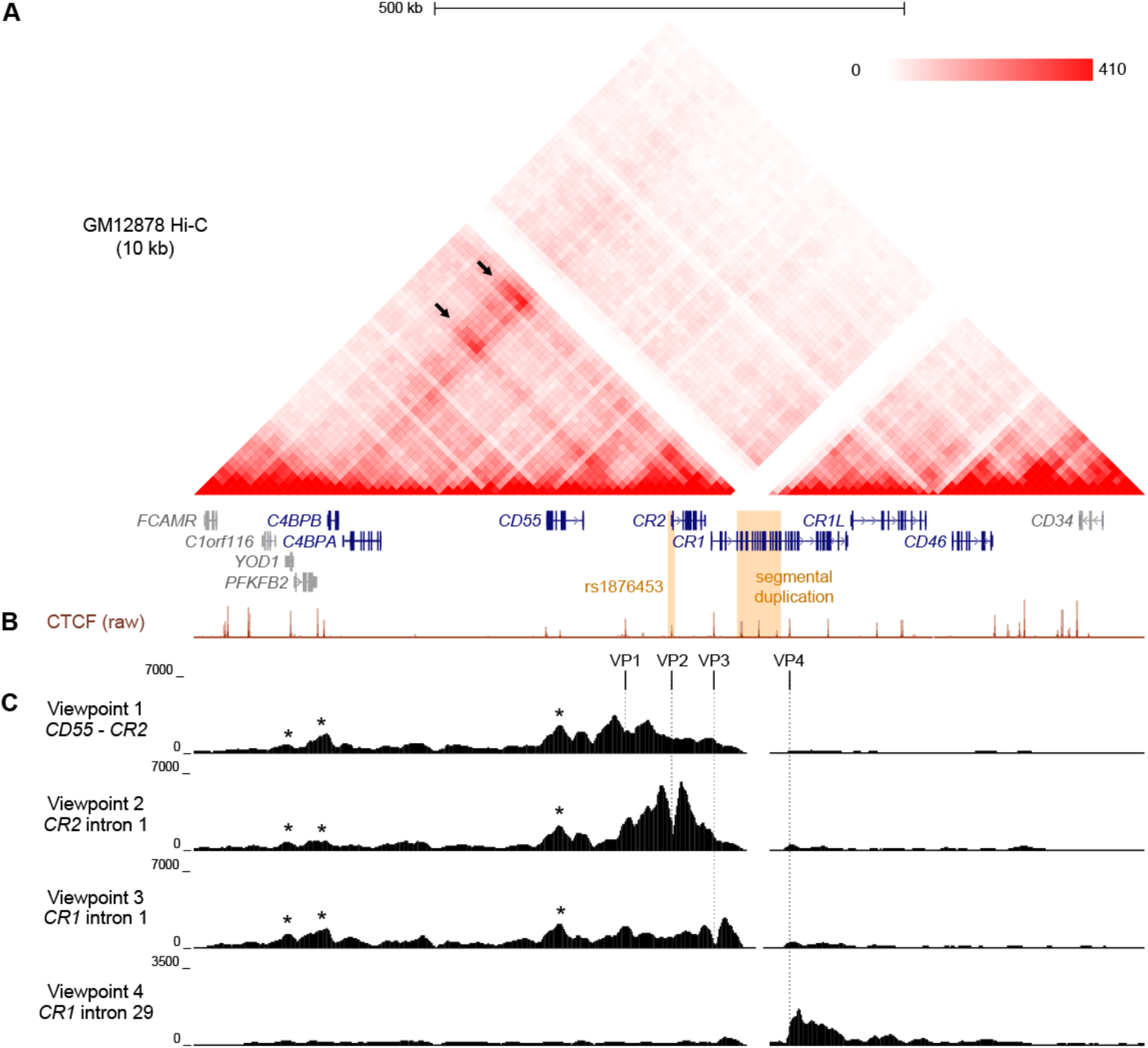
Chromatin conformation of the RCA gene cluster in B cells. A. Hi-C heatmap matrix (10 kb resolution) for the GM12878 B cell line from Rao *et al*. (41) for the 1 Mb region across the RCA genes (dark blue) on hg19 (chr1:207,120,000-208,130,000). Relative interaction frequencies between two loci are indicated by colour intensity (range 0-410). High frequency long-range interactions (>350 kb) were observed between distal RCA genes *C4BPB* and the complement receptor genes (*CR2* and *CR1*) (arrows), as well as between intervening loci, indicating that these genes reside in the same TAD. SLE-associated variants are indicated in orange. B. GM12878 ChIP-seq signal for CTCF from ENCODE shows CTCF enrichment at multiple sites across the RCA gene cluster which may engage in long-range chromatin looping. C. Chromatin conformation of the RCA gene cluster was fine-mapped using 4C-seq in the B-0028 cell line. Maps were generated from four viewpoints on CTCF binding sites in the intergenic region between *CR2* and *CD55* (viewpoint 1, VP1), intron 1 of *CR2* (viewpoint 2, VP2), the intron 1 of *CR1* (viewpoint 3, VP3) and intron 29 of CR1 (viewpoint 4, VP4). Viewpoints are represented by vertical dotted lines. Several 4C-seq peaks were common between VP1 – 3 and aligned with CTCF binding sites within *YOD1*, upstream of *C4BPB* and within *CD55* (asterisks). VP4 showed a distinct interaction profile to all other viewpoints.

To further clarify the TAD organisation of the RCA gene cluster, we performed 4C-seq in an analogous B lymphoblastoid cell line (B-0028). It is well established that in the B cell lineage, only membrane-bound RCA members (*CD55, CR2, CR1, CD46)* are expressed, not soluble protein members (*C4BPB* and *C4BPA*) (11). We confirmed this expression pattern using qPCR in the B-0028 cell line (**Supplementary Figure 2**). As CTCF plays an important role in chromatin looping, we selected 4C viewpoints (VP) from CTCF sites utilising B cell ChIP-seq data. (GM12878) Specifically viewpoints were selected which co-localised to regions of highly frequent interactions observed in the Hi-C data (**Figure 1B**). These viewpoints included; the intergenic region between *CD55* and *CR2* (VP1), intron 1 of *CR2* (VP2), which is the CTCF site influenced by SLE-associated SNP rs1876453 (*29*), and intron 1 of *CR1* (VP3) (**Figure 1B**). We also selected a 4C viewpoint from a CTCF binding site within intron 29 of *CR1* (VP4) which did not markedly engage in chromatin interactions with the upstream region of this gene cluster (**Figure 1B**). We confirmed enrichment of CTCF at these VPs in the B-0028 cell line using ChIP-qPCR (**Supplementary Figure 3**).

4C maps from VP1, VP2 and VP3 yielded consistent 4C signal peaks at upstream CTCF sites near RCA member *C4BPB* and within non-RCA member *YOD1* (**Figure 1C**, asterisks), corresponding to Hi-C data (**Figure 1A**). These CTCF viewpoints also consistently interacted with the CTCF site within intron 6 of RCA gene *CD55* (**Figure 1C**, asterisks). Chromatin interactions from VP1 – 3 did not extend to CTCF sites upstream of *YOD1* or downstream of *CR1* exon 7 (**Figure 1C**). In contrast, VP4 produced a unique 4C map whereby interactions were constrained to a 60 kb region downstream of this viewpoint within the *CR1* gene body and did not extend upstream (**Figure 1C**). We replicated 4C-seq from these CTCF viewpoints in another B cell line (B-0056), whereby CTCF interactions in the RCA gene cluster were also organised to two discrete regions (**Supplementary Figure 4**), potentially representing two TADs with an inter-TAD boundary located within the *CR1* gene itself. However, like Hi-C, reads mapping to the *CR1* segmental duplication were filtered out during 4C-seq data processing.

To refine the location of the TAD boundaries in the RCA gene cluster, we used a customized mHi-C pipeline which probabilistically assigns multi-mapping reads in Hi-C experiments to their most likely genomic position (42). Indeed, mHi-C successfully recovered Hi-C interactions across the *CR1* segmental duplication in the GM12878 cell line at 5 kb resolution (**Figure 2A**). These interactions were visualised as virtual 4C signal from RCA gene 5’ upstream promoter regions using Juicebox (43) (**Figure 2B**). In line with our previous observations, interactions from the promoters of upstream RCA genes *C4BPB, C4BPA, CD55, CR2* and *CR1* were localised to a distinct domain region compared to downstream RCA genes *CR1L* and *CD46* (**Figure 2B**). After recovering multi-mapping reads, we used spectralTAD (44) to systematically call TADs. Using this TAD caller, TAD boundaries were assigned at the intergenic region upstream of *YOD1*, intron 11 of *CR1* and the intergenic region downstream of *CD46*, placing RCA genes *C4BPB, C4BPA, CD55, CR2 and CR1* in TAD 1, and *CR1L* and *CD46* in TAD 2 (**Figure 2B**).

**Figure 2:**
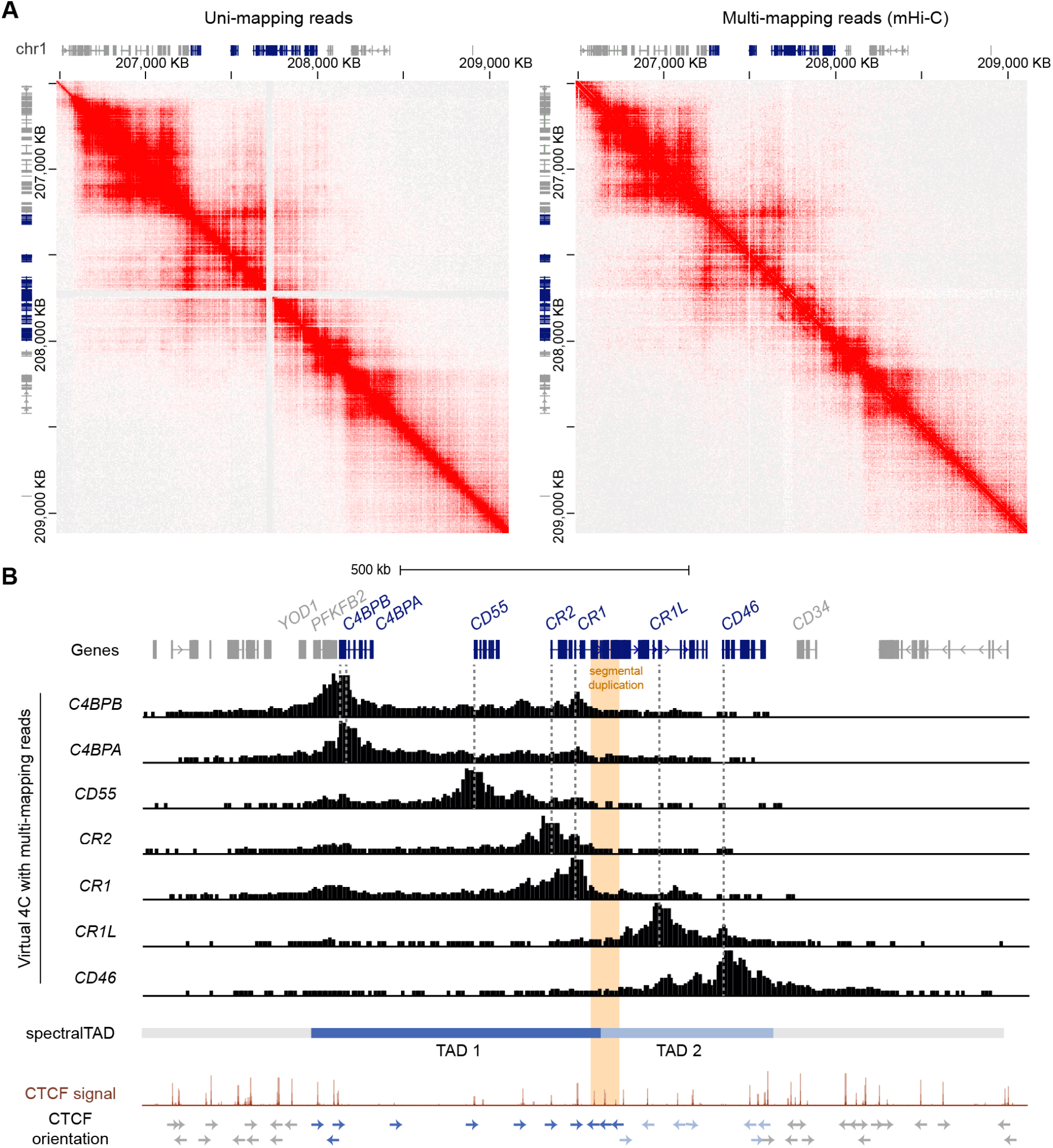
The RCA gene cluster is divided into two TADs and the inter-TAD boundary is located at the CR1 segmental duplication. A. Hi-C contact maps of GM12878 data at 5 kb resolution from Rao *et al*. (*44*) with only uni-mapping reads and both uni- (left panel) and multi-mapping (right panel) reads assigned with mHi-C. Using mHi-C, interactions across the *CR1* segmental duplication were successfully recovered. Data was visualised using Juicebox. B. mHi-C interactions were visualised as virtual 4C signal from the promoters of the RCA genes, which showed that interactions of RCA genes were constrained to two distinct regions. SpectralTAD was used to call TADs using Hi-C data with multi-mapping reads recovered, defining two clear TADs in the RCA cluster. Raw GM12878 CTCF ChIP-seq signal from ENCODE showed enrichment of convergent CTCF at the boundaries of both TADs and revealed that the *CR1* segmental duplication (orange) is flanked by repeated reverse orientation CTCF sites (indicated by arrows).

To corroborate these findings, we examined CTCF enrichment and motif binding orientation at the TAD boundaries in the RCA gene cluster. Each TAD was flanked by CTCF enrichment in the GM12878 B cell line and convergent CTCF motifs, characteristic of TAD boundaries. Importantly, the inter-TAD boundary was directly located to the *CR1* segmental duplication. Like other highly repetitive, unmappable genomic regions, CTCF enrichment at this region is underrepresented in the high-throughput datasets such as the ENCODE portal following the typical ChIP-seq processing pipeline. Therefore, we examined raw CTCF ChIP-seq signal in the GM12878 cell lines and observed enrichment of CTCF at this repeat element underlined by reverse orientation CTCF motifs at each repeat segment which has not been previously defined (**Figure 2B**). Enrichment of CTCF at the *CR1* repeat segments (*CR1* intron 7, 15 and 23) was observed across all cell lines in this ENCODE CTCF signal data set (**Supplementary Figure 5**). Taken together, our data shows that the RCA gene cluster is divided into two TADs and reveals that a TAD boundary is located within the *CR1* gene at the intragenic segmental duplication.

### Putative B cell enhancers in the RCA gene cluster were predicted to regulate multiple RCA genes

CTCF plays an important role in establishing long-range contacts within TADs and mediating enhancer-gene interactions (30). As CTCF-mediated chromatin looping was identified in the RCA gene cluster in B cells **(Figure 1**), we speculated that enhancer elements were present in this region that may regulate these genes in this cell lineage. To identify putative enhancers in the RCA, we leveraged a well-known enhancer database, GeneHancer. This integrates enhancer datasets from multiple consortium-based projects and other functional datasets to generate enhancer predictions and identify their potential gene-targets (45). Confidence scores for each enhancer prediction (GeneHancer score) and enhancer-gene prediction (gene-association score) were computationally assigned, based on the level of evidence retrieved. A strength of this database is that predicted enhancers can be classified as “double elite” if both their GeneHancer and gene-association scores were derived from more than one source of data, thus representing a prediction which is more likely to be functional (45). Numerous predicted enhancers on GeneHancer were identified across TAD 1 and TAD 2, but only a subset of these were classified as “double elite” (**Figure 3A, Figure 3B, Supplementary Table 4**).

**Figure 3:**
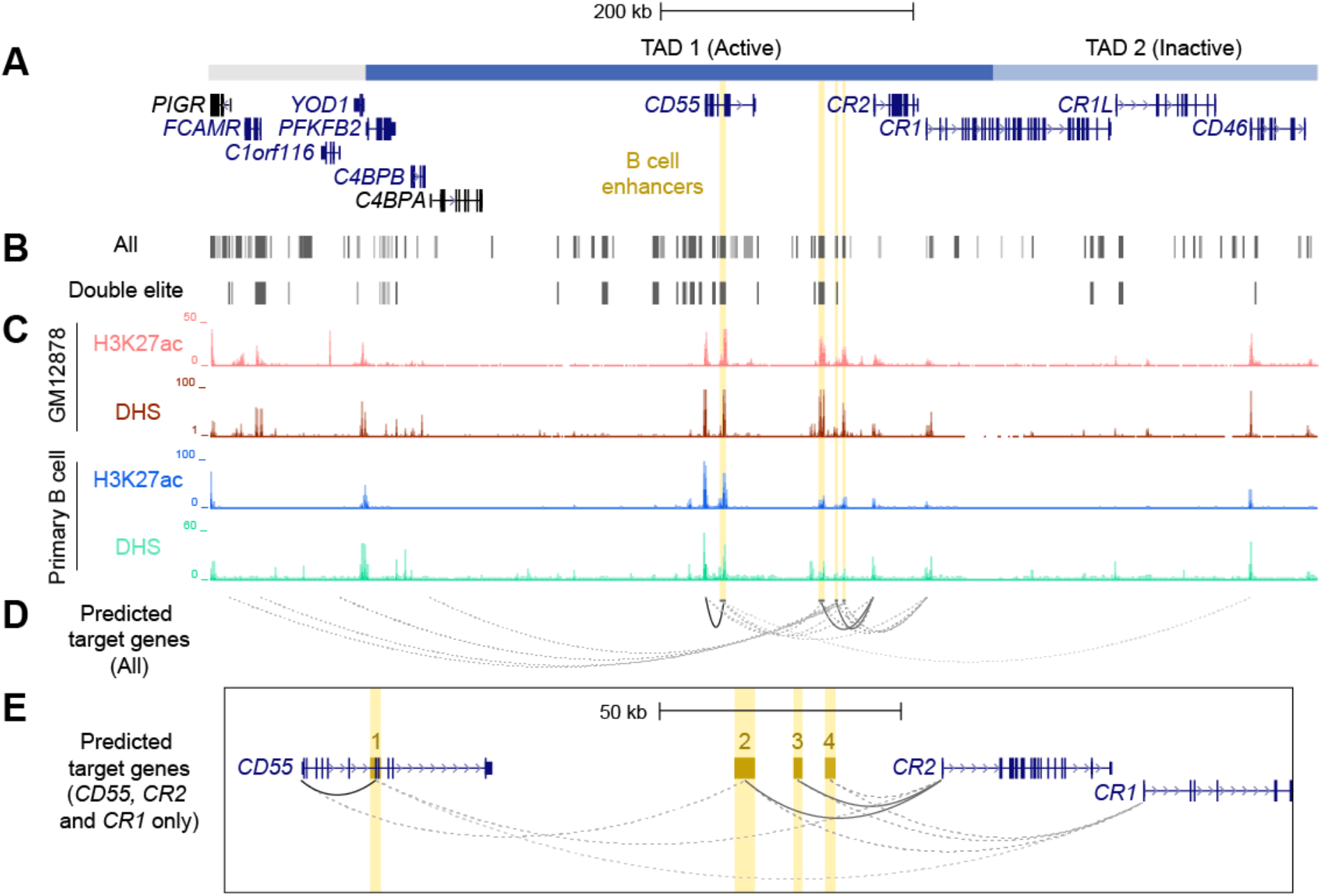
Identification and prioritisation of candidate B cell enhancers in TAD 1. A. The structural organisation of the RCA genes (dark blue) and upstream genes (grey) *PIGR, FCAMR, C1orf116* and *YOD1* on hg19 (chr1:207,104,491-207,978,031). B. Putative enhancers were identified by GeneHancer from multiple datasets from different consortia, such as ENCODE, Ensembl and FANTOM5 (GeneHancer). Each putative enhancer was also assigned predicted gene targets based on one or more methods. However, only a subset of putative enhancers were classified as ‘double elite’ on GeneHancer (Double elite). C. Four candidate B cell enhancers (yellow) were identified using ENCODE data for H3K27ac enrichment and DNase I hypersensitivity in different B cell samples (GM12878 B cell line and primary B cells from peripheral blood). D. Candidate B cell enhancers were predicted to regulate multiple genes. Target gene predictions that were identified by more than one method in GeneHancer are represented by a solid line. Predictions that were identified by just one method are represented by a dotted line. The opacity of each line represents the relative score/confidence for each gene-enhancer prediction as determined by GeneHancer whereby higher confidence predictions are darker. Scores determined by GeneHancer are listed in **Supplementary Table 5**. E. Region across *CD55, CR2* and *CR1* (exon 1 – 6) on hg19 (chr1:207,484,047-207,700,935). Candidate B cell enhancers (BEN) were named based on order of chromosomal position (BEN-1, BEN-2, BEN-3 and BEN-4). Evidence for BENs to regulate *CD55, CR2* and *CR1* was strongest among all gene-enhancer predictions.

Enhancers are important in cell-type specific regulation of gene expression and act by looping to their target gene promoters (46). To identify active enhancers that were most likely functional in B cells, we examined epigenetic marks characteristic of enhancers, such as H3K27ac and DNase I hypersensitivity (DHS) within candidate regions. We identified four candidate B cell enhancers (BENs) in TAD 1 that showed strong consistent H3K27ac enrichment and DHS in both B cell lines and primary B cells (**Figure 3C**). These candidate enhancers were located within *CD55* (BEN-1) or the intergenic region between *CD55* and *CR2* (BEN-2, BEN-3 and BEN-4) (**Table 1**). Furthermore, each candidate BEN contained binding sites for numerous transcription factors (based on ENCODE ChIP-seq data) including those important in B cell development, such as early B cell factor 1 (EBF1) (47) and PAX5 (48), and general regulatory factors (eg. EP300, CTCF and RNA polymerase II) (**Table 1**). The four BENs identified were supported by multiple lines of evidence to be active enhancer elements in B cells and were prioritised for further investigation.

**Table 1:** Candidate B cell enhancers (BENs) on GeneHancer were identified from multiple enhancer databases, contained numerous transcription factor binding sites (TFBS) and were predicted to regulate several genes.

|  | BEN-1 | BEN-2 | BEN-3 | BEN-4 |
| --- | --- | --- | --- | --- |
| GeneHancer ID | GH01J207333 | GH01J207411 | GH01J207424 | GH01J207429 |
| Location | *CD55*<br>(exon 5 – 8) | Intergenic<br>(*CD55* – *CR2*) | Intergenic<br>(*CD55* – *CR2*) | Intergenic<br>(*CD55* – *CR2*) |
| Enhancer sources | FANTOM5<br>ENCODE<br>Ensembl<br>dbSUPER | FANTOM5<br>ENCODE<br>Ensembl | FANTOM5<br>ENCODE<br>Ensembl | ENCODE |
| GeneHancer score | 1.6* | 1.2* | 1.1* | 0.7 |
| TFBS ^ | 64<br>EBF1 IRF4 PAX5<br>RELA SPI1 CTCF<br>EP300 POLR2A | 35<br>EBF1 IRF4 PAX5<br>RELA EP300<br>POLR2A | 13<br>EBF1 RELA SPI1<br>CTCF POLR2A | 22<br>EBF1 RELA SPI1 |
| Predicted gene targets | *CD55* * *CR2* *CR1*<br>*CD46* | *CR2* * *CR1* *CD55* | *CR2* * *CR1*<br>*C1orf116* *C4BPA* | *CR2* *CR1* *CD55*<br>*FCAMR* *PIGR* |

^ Only transcription factors important in B cell development (EBF1, IRF4, PAX5, RELA, SPI1) and gene regulation or chromatin organisation (CTCF, POL2RA and EP300) assayed using ChIP-seq in ENCODE are listed.
\* These predicted enhancers and gene-enhancer interactions were identified using more than one method by GeneHancer.

Predicted enhancers on GeneHancer were assigned putative gene targets using multiple methods and datasets, including expression quantitative trait loci (eQTL) analysis, enhancer-promoter interactions generated by capture Hi-C in the GM12878 cell line (CHi-C) and eRNA-mRNA co-expression from the FANTOM5 Enhancer Atlas (38, 45, 49). Each candidate BEN was predicted to regulate multiple genes, including RCA genes (*C4BPA, CD55, CR2, CR1* and *CD46*) and non-RCA genes (*PIGR, FCAMR, C1orf116*) (**Figure 3D, Supplementary Table 5**). However, only interactions between BEN-1 and *CD55*, BEN-2 and *CR2*, and BEN-3 and *CR2*, represented high-confidence (“elite”) associations, being identified by more than one contrasting method (**Figure 3E, Supplementary Table 5**). Of these, only BEN-1 was predicted to regulate a gene (*CD46*) located downstream of the intragenic TAD boundary in *CR1* (**Figure 3D**). However, this predicted interaction had the lowest score among all gene-enhancer predictions for these BENs (**Figure 3D, Supplementary Table 5**). Although these gene-enhancer interactions were based on bioinformatic predictions, this highlighted the potential for the RCA genes to be co-regulated in B cells.

### Candidate B cell enhancers in the RCA gene cluster were functional *in vitro*

To test the functionality of each BEN, we performed luciferase reporter gene assays using a constitutive minimal promoter (SV40) to drive luciferase expression. Each BEN was cloned upstream of the SV40 promoter in both forward and reverse orientation and the transcriptional effects were assayed in a panel of B cell lines (Reh, Raji, B-0028, SKW) and a non-B cell control (HepG2, liver cell-type) (**Figure 4**). Interestingly, transcriptional activity patterns of BEN-1 and BEN-3 were not consistent with that of an active enhancer, such that activity was unchanged or reduced relative to the control (pGL3-P, no enhancer) across the B cell lines, the latter indicative of silencer activity (**Figure 4**). BEN-4 displayed some enhancer activity in B cell lines but the relative increase in transcriptional activity was only significant in the SKW cell line in the reverse orientation (*p* = 0.0368, *n =* 3). (**Figure 4**). In contrast, BEN-2 significantly increased luciferase activity by approximately 3-fold relative to the control in SKW, in both forward (*p =* 0.0219, *n =* 3) and reverse orientation (*p =* 0.0436, *n =* 3), and by 1.5-fold in Raji in the forward orientation (*p =* 0.0003, *n =* 4) (**Figure 4**). Notably, transcriptional activity of BEN-2 was significantly decreased by 50% in the non-B cell line control (HepG2) in both enhancer orientations (forward *p =* 0.0321, *n =* 4; reverse *p =* 0.0255, *n =* 3) (**Figure 4**). Together, these data indicated that BEN-2 was the most likely candidate BEN to be active in the B cell lineage.

**Figure 4:**
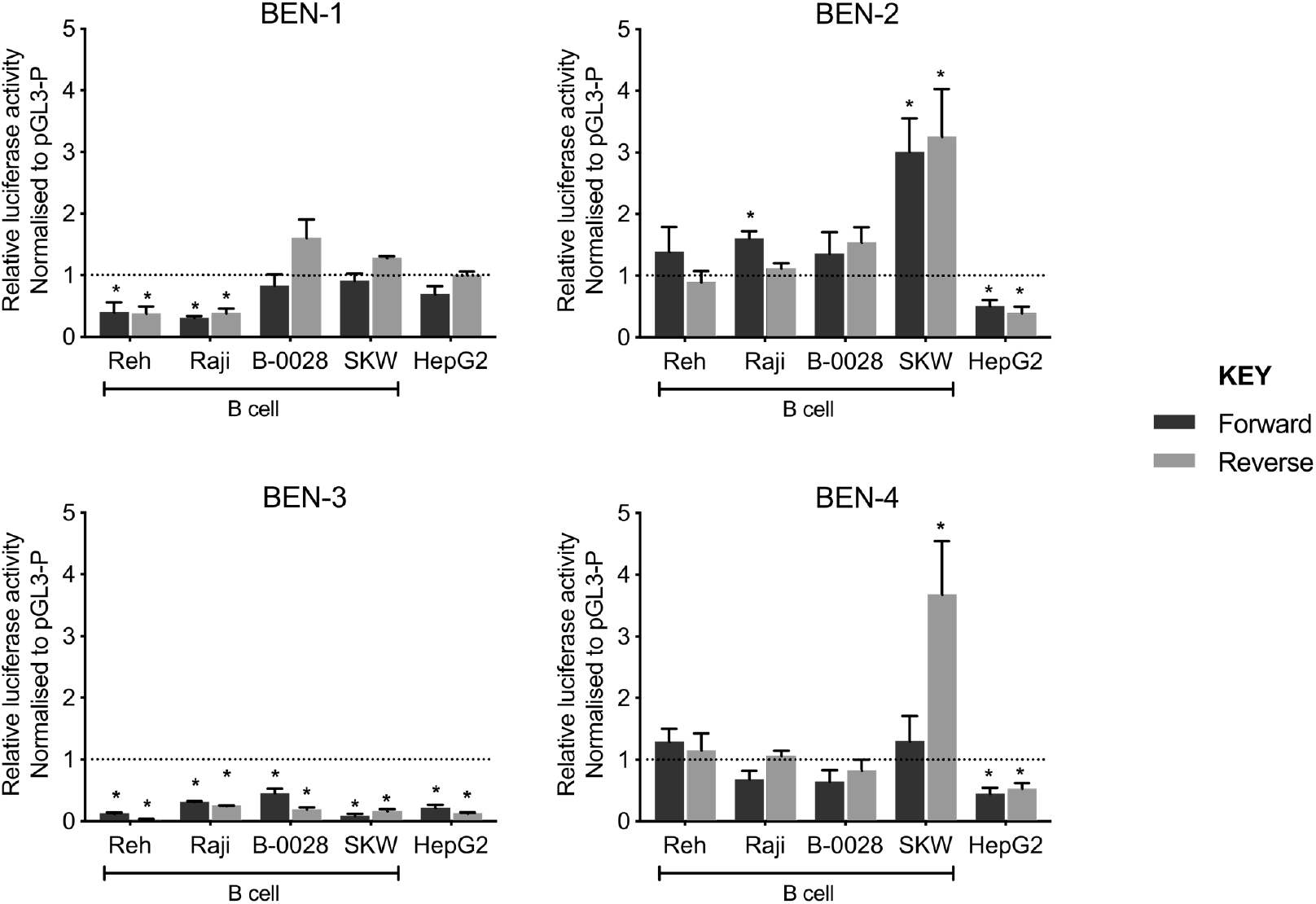
Candidate B cell enhancers demonstrated regulatory potential in luciferase assays, and BEN-2 increased relative transcriptional activity across a panel of B cell lines. Enhancer constructs for strong candidate B cell enhancers were cloned into the pGL3-P (Promega) luciferase plasmid, upstream of an SV40 minimal promoter in forward (black) and reverse (grey) orientation. Bars represent mean relative luciferase activity ± SEM after normalisation to an empty pGL3-P (no enhancer) control plasmid (*n =* 3 to 8). Asterisks represent statistically significant differences between normalised values and the pGL3-P control (*p* < 0.05). Dotted line at *y* = 1 represents normalised pGL3-P control value.

To support the functional role of BEN-2 in this cell type, we quantified epigenetic marks characteristic of enhancer regions, such as chromatin accessibility and H3K27ac enrichment, in the panel of cell lines used in the luciferase assays. Nucleosome occupancy was measured using MNase, a micrococcal nuclease which cannot bind and digest nucleosome-bound DNA. At BEN-2, nucleosome occupancy was consistently lower in the B cell lines than in the non-B cell control, HepG2 (**Figure 5A**). Conversely, chromatin accessibility was consistently high across all B cell lines, but inaccessible in HepG2 (**Figure 5A** and **Figure 5B**), indicating that this region is transcriptionally active in the B cell lineage. Accordingly, H3K27ac enrichment at BEN-2 was not observed in HepG2 but enriched in all B cell lines (**Figure 5C**). Together, these data are in support of BEN-2 acting as a functional B cell enhancer *in vitro*.

**Figure 5:**
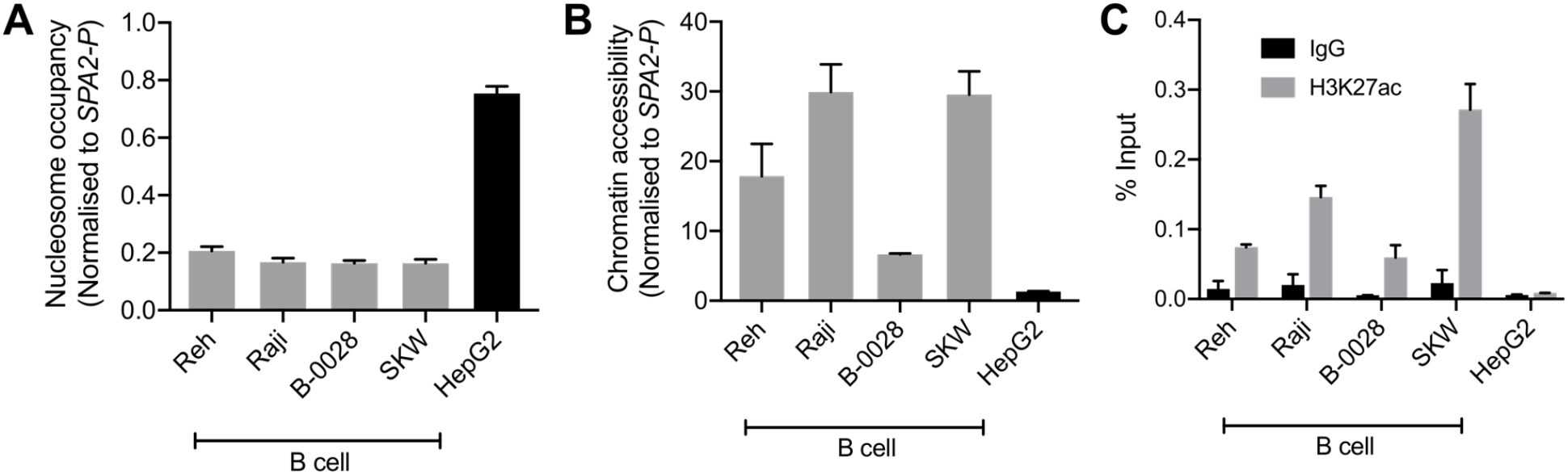
BEN-2 shows B cell-specific nucleosome occupancy, chromatin accessibility and enrichment for the H3K27ac active enhancer histone mark across a panel of B cell lines and non-B control (HepG2, liver). A. Nucleosome occupancy at BEN-2 as measured by ChART-PCR with MNase digestion. Data was normalised to the inaccessible *SFTPA2* gene promoter such that a value of 1.0 represents fully compacted nucleosomes, and lower values indicate less compacted nucleosomes. B. Chromatin accessibility at BEN-2 as measured by ChART-PCR with DNase I digestion. Data have been normalised to the inaccessible *SFTPA2* gene promoter. C. H3K27ac enrichment at BEN-2 as determined by ChIP-qPCR using the percent input method. Grey bars indicate H3K27ac enrichment at the target locus, and black bars show enrichment using a non-specific IgG control antibody. All data are presented as mean ± SEM from at least 3 biological replicates.

### CRISPR deletion of an intergenic B cell enhancer (BEN-2) decreased *CR2* and *CD55* expression at the transcript and protein levels

As reporter gene assays remove regulatory elements from their genomic context, which is an important aspect of enhancer function, we sought to assess the functional activity of BEN-2 *in vivo*. We also wished to confirm the predicted gene targets of BEN-2 identified on GeneHancer, including *CD55* and *CR2* which directly flank the enhancer (**Figure 3**). CRISPR deletion machinery was delivered using a plasmid-based method into the Raji mature B cell line. This cell line expresses *CD55, CR2* and *CD46*, although *CR1* is not expressed at levels detectable by qPCR (**Figure 6A**). This pattern of gene expression is in accordance with other B cell lines, such as B-0028 (**Supplementary Figure 2**). To efficiently delete the BEN-2 region, we modified the PX458 CRISPR plasmid to express two gRNA sequences that cut either side of BEN-2 (**Figure 6B**). The CRISPR plasmids, containing a GFP marker, were delivered into Raji cells and successfully transfected GFP-positive cells were enriched by fluorescence activated cell sorting (FACS). The resultant polyclonal GFP+ bulk population (bulk) was expanded and used for single cell cloning by limiting dilution. Successful enhancer deletion was qualitatively assessed using PCR (**Figure 6B, Figure 6C**). Indeed, PCR indicated that BEN-2 was deleted in a proportion of cells in the bulk population (**Figure 6C**). After screening expanded single cell clones, we successfully isolated a population containing a homozygous deletion of the BEN-2 region (**Figure 6C**). We confirmed the genotype of the del/del population using Sanger sequencing (**Supplementary Figure 6**).

**Figure 6:**
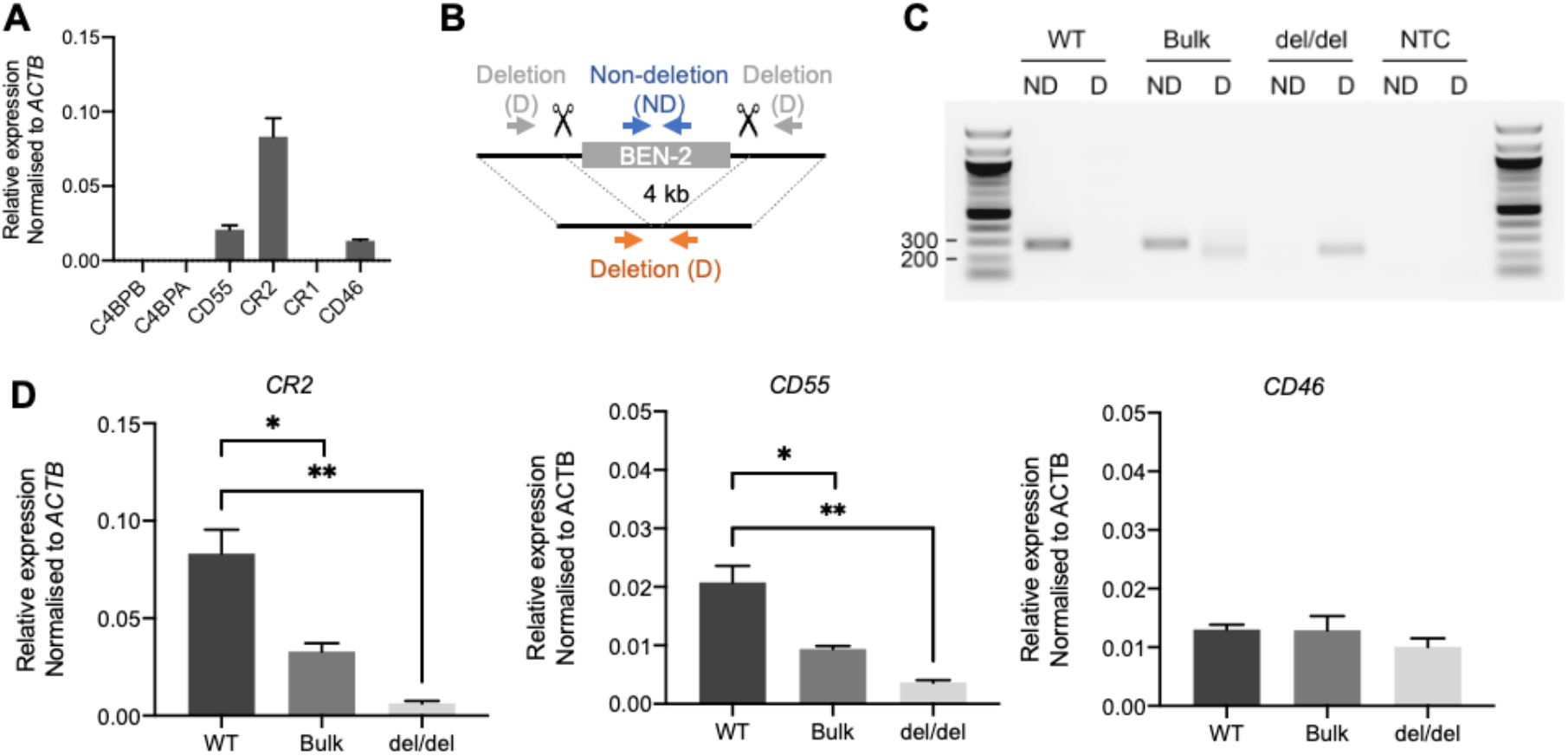
CRISPR deletion of BEN-2 decreased *CR2* and *CD55* mRNA expression in the Raji mature B cell line. A. Transcript abundance of RCA genes in Raji mature B cell line as measured by qPCR. Values were normalised to the β-actin gene (*ACTB*) using the ΔCt method. Bars represent mean relative expression ± SEM from at least 3 biological replicates. B. Schematic of enhancer deletion and screening strategy using CRISPR-Cas9. Deletion (4 kb) of BEN-2 was mediated by two gRNAs that cut either side of the enhancer region. Plasmids were modified from PX458 to express the two guides, Cas9 and a GFP marker. Screening was performed using PCR primers that flank the enhancer region (deletion; D) which amplify only in cases where a deletion has occurred (orange arrows). PCR primers that amplify within the enhancer region (non-deletion; ND) were used as a control (blue arrows). C. PCR deletion screen of wild-type Raji DNA (WT), polyclonal GFP+ bulk population (bulk) and monoclonal single-cell clone containing a homozygous deletion of BEN-2 (del/del). PCRs were run alongside a no-template control (NTC). D. Transcript abundance of RCA genes in TAD 1 (*CR2* and *CD55*) and TAD 2 (*CD46*) were measured by qPCR. Values were normalised to the β-actin gene (*ACTB*) using the ΔCt method. Bars represent mean relative expression ± SEM from 3 biological replicates. Asterisks represent statistically significant differences between WT, bulk and del/del samples (\**p* < 0.05, \*\**p* < 0.005).

Remarkably, the homozygous deletion of BEN-2 in Raji cells significantly decreased *CR2* transcript abundance by approximately 90% relative to WT levels (*p =* 0.0034, *n =* 3) and *CD55* transcript abundance by approximately 80% of WT levels (*p* = 0.0039, *n = 3*). We also measured transcript abundance of *CR2* and *CD55* in the bulk population where a proportion of cells contained the enhancer deletion. Accordingly, both transcripts were significantly decreased compared to WT but to a lesser extent than del/del population; *CR2* transcript abundance decreased by approximately 70% of WT levels (*p =* 0.0184, *n =* 3) and *CD55* by approximately 50% of WT levels (*p =* 0.0167, *n =* 3) (**Figure 6D**). We also measured transcript abundance of *CD46*, which was not predicted to be targeted by BEN-2 on GeneHancer and localised to the neighbouring TAD (**Figure 3**). Enhancer deletion did not alter *CD46* transcript abundance (**Figure 6C**). These data confirmed that BEN-2 is a functional enhancer in B cells and demonstrated that BEN-2 regulates *CD55* and *CR2* within this cellular context.

As *CR2* and *CD55* transcript levels were significantly decreased with BEN-2 deletion in the Raji cell line, we next determined if surface protein expression of these receptors was concomitantly affected. We used flow cytometry to assess CR2 surface expression in the bulk and del/del populations. Indeed, CR2 expression was significantly decreased by approximately 3-fold in del/del population relative to the WT control (*p =* 0.0257, *n =* 3) (**Figure 7A, Figure 7B**). Surface staining using a CD55 antibody showed that around 2-3% of the population of Raji cells were CD55-positive (**Figure 7B, Figure 7C**). Regardless, the overall CD55 surface expression was also significantly decreased to approximately 75% of WT levels (*p =* 0.074, *n =* 3) (**Figure 7A**). In line with transcript abundance data (**Figure 6D**), CR2 and CD55 surface expression in the bulk population was decreased to levels between WT and del/del; 72% and 90%, respectively, of WT levels. This confirmed that the reduction in CR2 and CD55 transcript expression with CRISPR deletion of BEN-2 reduced surface protein levels. We thus further validated that BEN-2 regulates these genes both at the level of mRNA and subsequent protein expression in a B cell context.

**Figure 7:**
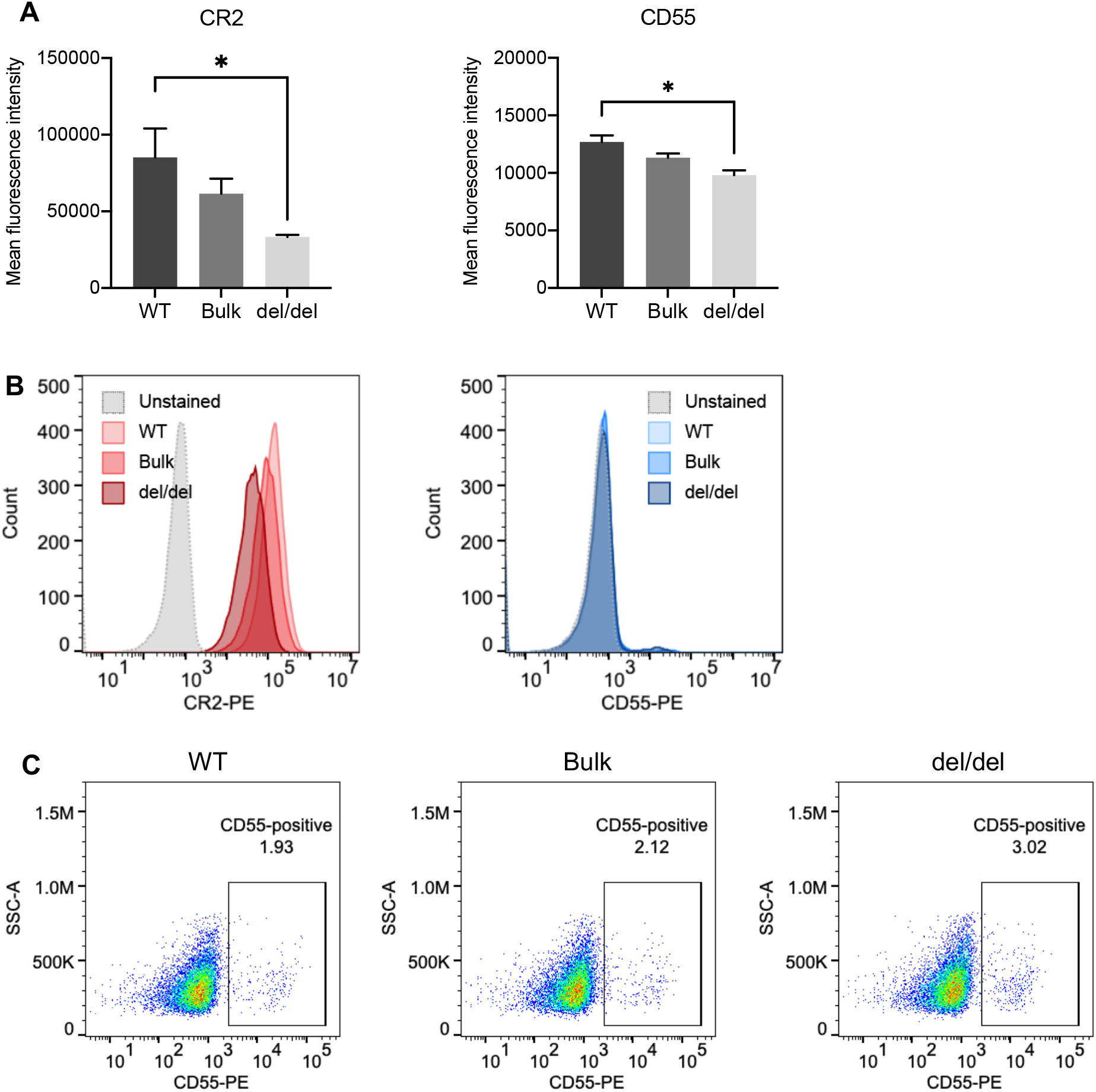
CRISPR deletion of BEN-2 decreased surface expression of CR2 and CD55 in the Raji B cell line. A. Cell surface expression of CR2 and CD55 protein was determined using flow cytometry. Cells were labelled with PE-conjugated CR2 antibody, PE-conjugated CD55 antibody or PE-conjugated IgG (isotype control) to confirm positive expression. Samples were run alongside unstained controls (not shown). For each sample, 10000 events were collected. Bars represent mean fluorescence intensity ± SEM from 3 biological replicates. Asterisks represent statistically significant differences between WT and del/del samples (*p* < 0.05). B. Representative histograms from the WT, bulk and del/del samples stained with CR2-PE or CD55-PE, as well as unstained WT control. C. Representative dot plot from the WT, bulk and del/del samples stained with CD55-PE stained samples. The CD55-positive gate was plotted using the unstained control.

## DISCUSSION

In this manuscript, we have explored the chromatin architecture of the RCA gene cluster, an important immunomodulatory region. We provide multiple insights into genes and variants within this cluster from a biological, evolutionary and disease perspective. We show for the first time the co-localisation of the RCA genes is in part due to the requirement of shared long-range regulatory elements. Using high-resolution 4C-seq maps in B cell models, we showed that several distal CTCF sites in the RCA gene cluster engage in chromatin looping, including the CTCF site modulated by the SLE-associated SNP rs1876453 (29). We further shine a light onto the enigma of the unmappable segmental duplication within *CR1* (24). Overall, we reveal extensive and complex mechanisms by which the RCA gene cluster may influence gene expression and thereby autoimmunity.

While Hi-C interaction data are typically sufficient to identify TADs across the genome, the utility of these data to examine the structural arrangement of the RCA gene cluster was hindered by the sequence repetitiveness of the intragenic segmental duplication in *CR1*. Our strategy combined high-resolution 4C-seq and a unique Hi-C pipeline (mHi-C) in conjunction with publicly available Hi-C datasets to uncover a highly intriguing TAD arrangement within the RCA gene cluster in the B cell lineage. We successfully utilised 4C-seq in two representative B cell lines from multiple viewpoints. Further, we were able to show consistent maps of CTCF interactions in the RCA gene cluster and chromatin looping which was constrained to one of two distinct regions across these loci. Importantly, when we examined two CTCF viewpoints within intron 1 and intron 23 of the *CR1* gene, we found that interactions from these viewpoints were directionally constrained in opposite directions (upstream and downstream of *CR1*, respectively). These data indicated that the inter-TAD boundary was located within the *CR1* gene. We corroborated this finding using mHi-C, a novel Hi-C processing pipeline which assigns multi-mapping reads to the most likely genomic location. We thus resolved interactions at this previously unmappable element and refined the location of the inter-TAD boundary to intron 11 of *CR1*, falling within the segmental duplication. Previous studies have shown that TAD boundaries are enriched within housekeeping genes (31, 50) and may also be located near gene promoters (51). To our knowledge, a TAD boundary located well within the body of an expressed protein-coding gene has not been delineated to this resolution prior to our study.

The tandem segmental duplication in *CR1*, also known as ‘low copy repeat 1’ (LCR1), results in the duplication of 8 exons and introns in *CR1* and has been shown to alter the number of functional domains in the protein (52, 53). There are multiple co-dominant *CR1* alleles in the population defined by copies of LCR1. CR1-A/F (one copy of LCR1) and CR1-B/S (two copies of LCR1) alleles are most common. Alleles that contain zero copies and three copies of LCR1 have also been documented (54). Importantly, this repeat element is known to be associated with SLE and, more recently, late-onset Alzheimer’s disease (24, 55). Despite the large size and nature of this repeat element, its biological implication is undefined. Our findings strongly indicate that the LCR1 repeat element in *CR1* co-localises with a TAD boundary and thus has a role in regulating gene expression in the RCA cluster. This is consistent with the presiding hypothesis that complex diseases develop as a result of dysregulated gene expression rather than functional protein changes. Based on the loop extrusion model of TAD formation, it is likely that the increasing copy number of LCR1 results in increased numbers of CTCF sites (reverse orientation) in TAD 1, thereby increasing insulation of chromatin interactions within this TAD (32). Experiments are currently on-going in our laboratory to establish whether the LCR1 copy number influences chromatin interactions and gene expression in the RCA cluster.

The RCA gene cluster is an exemplar gene cluster as its members are co-localised in the human genome and share protein structure and function. In addition, the genes, gene orientation and gene order of the RCA cluster are well conserved in many species across evolutionary time, including *Xenopus tropicalis* and chicken (56, 57). The RCA gene cluster in mice is also conserved but its members are separated across two chromosomal positions located more than 6 Mb apart (58). This matches closely to the TAD organisation of the gene cluster we describe here. It has been observed that breaks in synteny between species commonly occur at TAD boundaries (59), thus the TAD boundary we identified in *CR1* may represent the breakpoint region for the genomic rearrangement of the RCA gene cluster in humans and mice.

Our study revealed that the RCA gene cluster is divided into two TADs in the B cell lineage; TAD 1 consists of *C4BPB, C4BPA, CD55, CR2* and *CR1*, and TAD 2 consists of *CR1L* and *CD46*. This grouping does not reflect the organisation of active RCA genes in B cells (*C4BPB* and *C4BPA* are not expressed in B cells) or other recognised cell types. Nonetheless, our findings in the RCA gene cluster correspond to the *Six* (60), *HoxA* (61, 62) and *HoxD* (63, 64) homeobox gene clusters which have also been shown to be separated into two distinct TADs and regulatory regions. Unlike the aforementioned gene clusters, which have highly-restricted and distinct expression patterns (65), the RCA gene cluster is unique in that its members are expressed in numerous cell types and across various stages of cell development. Each member of the RCA has a distinct expression pattern; *CD55* and *CD46* are expressed across nearly all cell types, including non-immune cells (11), whereas *CR2* and *CR1* are predominantly expressed on B cells (66) and erythrocytes (67), respectively. Thus it will be interesting to examine the RCA gene cluster in other cellular contexts to examine the TAD structure and enhancer-gene landscape and determine if these differ from the organisation we define in this study. This will require multiple strategies of experimentation, as we have utilised here, to overcome the caveats of reliance on high-throughput datasets.

Gene-gene interactions are a long-recognised and important contributor to complex disease susceptibility, often overlooked due to the difficulty in identifying such interactions (68). In this study, we mapped such interactions between genes in the RCA gene cluster at the molecular level using high-resolution chromatin interaction maps and have successfully identified direct chromatin interactions between CTCF sites in *CD55, CR2* and *CR1*. Binding of CTCF at intron 1 of *CR2* was modulated by an SLE-associated SNP (rs1876453) and shown to influence the expression of its neighbouring gene, *CR1*, in B cells (29). Our data now shows that *CR2* and *CR1* form part of the same CTCF-mediated chromatin network in the RCA gene cluster and support the hypothesis that these genes are co-regulated. It is possible that expression of *CR1* is also co-regulated by BEN-2 in TAD 1 as has been predicted by CHi-C in the GM12878 B cell line (49).

We also uncovered a direct relationship between *CR2* and *CD55*, showing that these genes are co-regulated by an intergenic enhancer, BEN-2. This marks the very first long-range regulatory element identified in this TAD and gene cluster. We characterised this enhancer through multiple lines of evidence *in vitro* and *in vivo* in several B cell models; by characterising transcriptional activity and chromatin marks, as well as utilising CRISPR genomic deletion. Importantly, the reduction in transcript level mediated by this enhancer deletion produced concomitant reduction in surface protein level of both CR2 and CD55. This highlights the importance of BEN-2 in the expression of these complement regulators and improves our understanding of the complex transcriptional control of these genes.

The surface expression of CD55 (69, 70) and CR2 (19, 20) on B cells is significantly decreased in SLE patients. Genetic variation, such as SNPs, in the BEN-2 region may influence expression of both *CR2* and *CD55* in tandem, thereby exacerbating the effect of variants in their contributions to autoimmunity. The genes of the RCA cluster may also be co-regulated by the other candidate B cell enhancers we identified here. Strategies to map gene-gene interactions and define relationships between genes such as those utilised in this study may open important avenues to better understand how complex diseases, like SLE and Alzheimer’s, are influenced by genetic variation in this region.

We have established for the first time that the RCA gene cluster is transcriptionally co-regulated and comprises of a complex network of enhancer-gene and gene-gene interactions. We have also defined the regulatory architecture of the RCA in the B cell lineage, revealing novel mechanisms by which the RCA gene cluster is controlled and expanding the scope for future investigations in the context of evolution, immunity and complex genetic disease.

## MATERIAL AND METHODS

### Cell culture

Cell lines Reh (CRL-8286), Raji (CCL-86), SKW (TIB-215), K562 (CCL-243) and HepG2 (HB-8065), were obtained from the American Type Culture Collection. B lymphoblastoid cell lines (B-0028 and B-0056) were derived from healthy individuals and immortalised by Epstein-Barr virus infection (29). All suspension cells were cultured in RPMI-1640 with L-glutamine (Life Technologies), supplemented with 10% FBS, 100 μg/mL penicillin and 100 ng/μL streptomycin. The adherent cell line (HepG2) was cultured in high glucose DMEM (Life Technologies) with 10% FBS, 100 μg/mL penicillin and 100 ng/μL streptomycin.

### Circular chromosome conformation capture (4C-seq)

B-lymphoblastoid cell lines (5 × 10^6^ cells) were harvested by centrifugation and resuspended in 5 mL PBS with 10% FBS. To cross-link cells, 5 mL 4% formaldehyde was added, and samples were incubated for 10 min. Cross-linking was quenched by adding 1 M glycine to 125 mM final concentration and cells collected by centrifugation at 300 x *g* for 10 min at 4°C. 4C-seq assays and data processing were performed as previously reported (71, 72). Sequences of primers used as 4C viewpoints are listed in **Supplementary Table 1**.

### Bioinformatic datasets and pipelines

Hi-C data for GM12878 from Rao et al. (41) were visualised as contact heatmaps and virtual 4C signal using the 3D Genome Browser (73) and Juicebox (43). CTCF orientation calls from GM12878 were retrieved from Rao *et al*. (41) and assessed in the *CR1* segmental duplication using CTCFBSDB 2.0 (74). Enhancer predictions were retrieved from the GeneHancer database (Version J), which leverages data from multiple sources, including ENCODE, FANTOM5 and Ensembl. Histone modifications and transcription factor enrichment was assessed using ENCODE data and visualised on the UCSC Genome Browser on hg19.

### Mapping Hi-C reads with mHi-C

Multi-mapping Hi-C sequencing reads from Rao *et al*. (41) were evaluated using the mHi-C pipeline (42) at 5 kb resolution (**Supplementary Table 2**). mHi-C was used as described in Zheng *et al*. (42) with a novel post-mHiC processing strategy. In brief, the genomic distance effect on the contact probabilities is estimated using the univariate spline model based on uniquely mapping reads. Such prior probabilities information is updated iteratively by the local bin-pairs contact counts leveraging both uniquely mapping reads and multi-mapping reads. The posterior probabilities, as the results of mHi-C, quantify the chance for the candidate bin-pair to be the true origin for each multi-mapping read pair. Instead of applying general filtering based on the posterior score by a fixed threshold, the posterior probabilities are interpreted as fractional Hi-C contact counts to incorporate a more significant number of the multi-mapping reads into the analysis. To examine interaction artefacts due to highly repetitive sequences, a stringent multi-mapping allocation strategy was employed which enforced all the multi-mapping reads assigned to corresponding regions have greater or equal to a 0.99 posterior score. Subsequent to rescuing multi-mapping reads by mHi-C, TAD boundaries are detected by the state-of-the-art TAD caller spectralTAD (44) which provides nested TAD at different levels. The TADs shown in Figure 2 are first-level TAD boundaries called at 25 kb resolution.

### Luciferase reporter-gene assays

Candidate enhancers were amplified from human genomic DNA using Q5 Hot-Start High-Fidelity DNA polymerase (New England Biolabs) and directionally cloned into the pGL3-Promoter plasmid (pGL3-P) (Promega) upstream of the SV40 promoter using restriction enzymes. Plasmid DNA was prepared using the EndoFree Plasmid Maxi Kit (QIAGEN) for transfection. Each enhancer construct (1 μg) was transiently transfected with the pRL-TK *Renilla* internal control vector (50 ng) using 4 μL Viafect™ transfection reagent (Promega) into suspension cell lines or adherent cell lines. Cell lysates were harvested after 24 h of incubation. Firefly and *Renilla* luciferase activity of cell lysates were sequentially assayed using the Dual-Luciferase Reporter Assay System (Promega) on a GloMax Explorer luminometer (Promega). Firefly luciferase readings were normalised to a co-transfected internal *Renilla* luciferase control, and the activity of each enhancer construct was normalised to a pGL3-P control. Sequences of primers used in this paper are listed in **Supplementary Table 1**.

### Quantitative PCR

Total RNA was extracted from cells using the RNeasy Mini Kit (QIAGEN) with on-column DNase I treatment. RNA quantity and purity were determined by spectrophotometry. RNA was reverse-transcribed into cDNA using SuperScript III VILO reverse transcriptase (Life Technologies) and diluted with UltraPure dH_2_O (Life Technologies). qPCR reactions comprised 1X SYBR Green No-Rox (Bioline), 250 nM forward and reverse primers (**Supplementary Table 1**), and 2 μL diluted cDNA up to a final volume of 10 μL. Cycling and analysis were conducted using a Mic qPCR Cycler (BioMolecular Systems) with the following conditions: 95°C for 10 min, and 35 cycles of 95°C for 15 s, 60°C for 15 s, and 72°C for 15 s. Melt curve analysis was used to confirm specific amplification of targets. Relative mRNA expression levels were calculated using the comparative Ct method, normalised to β-actin (*ACTB*).

### Chromatin immunoprecipitation

Briefly, 4 × 10^7^ cells were fixed using 1% formaldehyde (Sigma-Aldrich) for 10 min. Cells were washed in PBS and lysed using NP-40 lysis buffer. Cell nuclei were resuspended in 2 mL 0.4% SDS shearing buffer for sonication with a Covaris S220X sonicator (Covaris) for 7 min. For each immunoprecipitation, 25 μg chromatin was diluted with IP dilution buffer and pre-cleared with Protein A agarose beads (Merck-Millipore) for 1 h at 4°C. Chromatin was incubated with 5 μL anti-CTCF (Merck-Millipore), 5 μg anti-H3K27ac (Abcam), or 5 μg rabbit IgG isotype control antibody (Merck-Millipore) for 16 h at 4°C with rotation. Immune complexes were collected by centrifugation and cleared using Protein A agarose beads (Millipore) and incubated for 1.5 h at 4°C. Complexes were washed and eluted in 500 μL ChIP elution buffer. Crosslinks were reversed by adding 25 μL 4M NaCl and incubation for 16 h at 65°C with shaking (600 rpm). Samples were treated with RNase A and Proteinase K, and DNA was purified using the QIAquick PCR Purification kit (QIAGEN) according to the manufacturer’s specifications using 50 μL Buffer EB. For analysis, 2 μL of purified DNA was used for qPCR reactions with a Mic qPCR cycler as described above. Enrichment was determined using the percent input method.

### Chromatin accessibility by real-time PCR

Chromatin accessibility by real-time PCR (ChART-PCR) was performed as previously described (75) using 20 U DNase I (Promega). To assess nucleosome occupancy, 1000 Gel Units MNase (New England Biolabs) was used. Digested and undigested samples were purified using the QIAquick PCR Purification kit (QIAGEN). For analysis, qPCR reactions consisting of 50 ng DNA, 1X SYBR Green (Bioline), 250 nM primers up to a final volume of 10 μL were cycled using a ViiA7 real-time thermocycler and QuantStudio V1.3 (Applied Biosystems). Cycling conditions were as follows: 95°C for 10 min, 40 cycles of: 95°C for 15 s, 60°C for 15 s, 72°C for 30 s, followed by melt curve analysis. Accessibility levels were determined using the comparative Ct method for undigested and digested samples, normalised to the lung-specific *SFTPA2* promoter (SPA2-P) control locus. For MNase nucleosome occupancy assays, normalised data were transformed such that a value of 1.0 represents completely compacted nucleosomes, and lower values indicate reduced nucleosome occupancy.

### CRISPR deletion

CRISPR plasmid constructs were modified from pSpCas9(BB)-2A-GFP (PX458), a gift from Feng Zhang (76) (Addgene plasmid #48138). To generate a large genomic deletion, PX458 was modified to express two guide RNAs (gRNAs) to cut the 5’ and 3’ ends of the target region. gRNAs were designed using CRISPRscan (77) to select highest scoring sequences with minimal off-target effects (**Supplementary Table 3**). gRNAs were cloned into the *Bbs*I restriction sites of PX458 using T4 DNA ligase (New England Biolabs). gRNA expression cassette inserts (U6 RNA polymerase III, gRNA sequence and gRNA scaffold) were amplified using PCR with primers containing oligonucleotides with *Acc*65I and *Xba*I restriction ends for sub-cloning (**Supplementary Table 1**). For the negative control construct, the gRNA expression cassette of PX458 was removed using *Pci*I/*Xba*I digestion and purified using the QIAquick Gel Extraction kit (QIAGEN). The linearised plasmid was blunted using T4 DNA polymerase (New England Biolabs) and re-ligated.

CRISPR plasmid constructs (2 μg) were electroporated into 2 × 10^6^ Raji cells using the Amaxa Cell Line Nucleofector Kit V (Lonza Bioscience) (Program M-013). Cells were incubated at 37°C with 5% CO_2_ for 24 h and then sorted for GFP+ expression using fluorescent activated cell sorting (FACS) on a FACSAria II (BD Bioscience). The GFP+ pool (bulk) was expanded for further analysis. To obtain single cell clones, the bulk sample was plated into 96-well plates at approximately 1 cell per well in conditioned media. Single cell colonies were expanded and cells were cryopreserved. Genomic DNA and total RNA extraction was performed using the QIAamp DNA Blood Mini kit (QIAGEN) and RNeasy Mini kit (QIAGEN), respectively. RNA was reverse-transcribed and transcript abundance was measured by qPCR as previously described.

### CRISPR deletion screening

DNA was qualitatively screened for the genomic deletion using PCR with oligonucleotides amplifying across the targeted region (deletion; D) and within the target region (non-deletion; ND) (**Supplementary Table 1**). DNA (50 ng) was amplified using 1X GoTaq Green Master Mix (Promega), 0.8 μM oligonucleotides and 5% DMSO up to a volume of 20 μL, and cycled as follows: 95°C for 5 min, followed by 30 cycles of 95°C for 30 s, 61°C for 18 s, 72°C for 5 s, and a final extension at 72°C for 5 min. Single cell clones that screened positive for genomic deletion were confirmed using Sanger sequencing. The deletion product from the bulk sample was also analysed using Sanger sequencing.

### Flow cytometry

Cells (1 × 10^6^ cells) were harvested and washed with cold staining buffer (PBS with 5% FBS (v/v)) at 300 × *g* for 5 min at 4°C. For surface staining, cells were resuspended in 90 μL staining buffer and incubated with 10 μL of anti-human CD21-PE (Cat #555422, BD Bioscience), anti-human CD55-PE (MHCD5504, Thermo Fisher Scientific) or IgG1κ-PE isotype control (Cat #555749, BD Bioscience) for 20 min on ice. After incubation, cells were washed and resuspended in 0.5 mL staining buffer and processed using a BD Accuri C6 flow cytometer (BD Bioscience). Data was analysed using FlowJo software V10.8.0 (Tree Star). Samples were run alongside unstained controls.

### Statistical analysis

Differences in transcriptional activity, mRNA expression and mean fluorescence intensity were assessed using Student’s unpaired t-test with a confidence interval of 95% (*p* < 0.05). Statistics and graphs were generated using GraphPad Prism version 7.0 (GraphPad). Graphed values represent the mean ± SEM of at least three independent experiments.

## ACCESSION NUMBERS

4C-seq data were deposited in the Gene Expression Omnibus (GEO) database under accession number GSE140127. The mHi-C pipeline can be accessed at https://github.com/yezhengSTAT/mHiC.

## ACKNOWLEDGEMENTS

First and foremost, we would like to thank and remember our late collaborator, J.L., for his integral and generous scientific guidance in this project, and express our deepest condolences to his family, friends and colleagues. We would like to thank Kevin Li and the FACS Facility at the Harry Perkins Institute of Medical Research for technical assistance with cell sorting. Further, we thank Kathy Fuller and Henry Hui for their technical expertise and reagents for flow cytometry.

## FUNDING

This work was supported by the National Institutes of Health [R01 AI24717 to J.B.H.], the Australian Government Research Training Program Scholarship at the University of Western Australia [to J.C. and J.S.C.], the Spanish Government [BFU2016-74961-P to J.L.G-.S] and an institutional grant Unidad de Excelencia María de Maeztu [MDM-206-0687 to the Department of Gene Regulation and Morphogenesis, Centro Andaluz de Biología del Desarrollo].

## CONTRIBUTIONS

J.C., J.S.C., R.D.A. and Y.Z. performed the experiments and conducted the bioinformatic analyses; J.C., J.S.C., J.L.G.-S., R.L.T. and D.U. designed the experiments; J.C., J.S.C., Y.Z., E.Q. and D.U. analysed results; J.S.C. and D.U. conceptualised the project; J.L.G.-S., R.L.T., S.K. M.F. S.A.B. and J.B.H. provided intellectual input and resources; E.Q. and D.U. supervised the project; J.C. wrote the manuscript draft with input from all authors. All authors reviewed the manuscript.

## CONFLICT OF INTEREST

The authors declare no competing interests.

## Notes

### Competing Interest Statement

The authors have declared no competing interest.

### Summary of Updates

Additional data added (Figure 2) to support evidence and identification of TAD organisation of the RCA cluster with new collaborators/authors; data also added (Figure 6 and Figure 7) to clarify the effect of the enhancer deletion on CR2 and CD55 expression.

