## Supplementary Figures and Tables for "Regulatory architecture of the RCA gene cluster captures an intragenic TAD boundary, CTCF-mediated chromatin looping and a long-range intergenic enhancer"

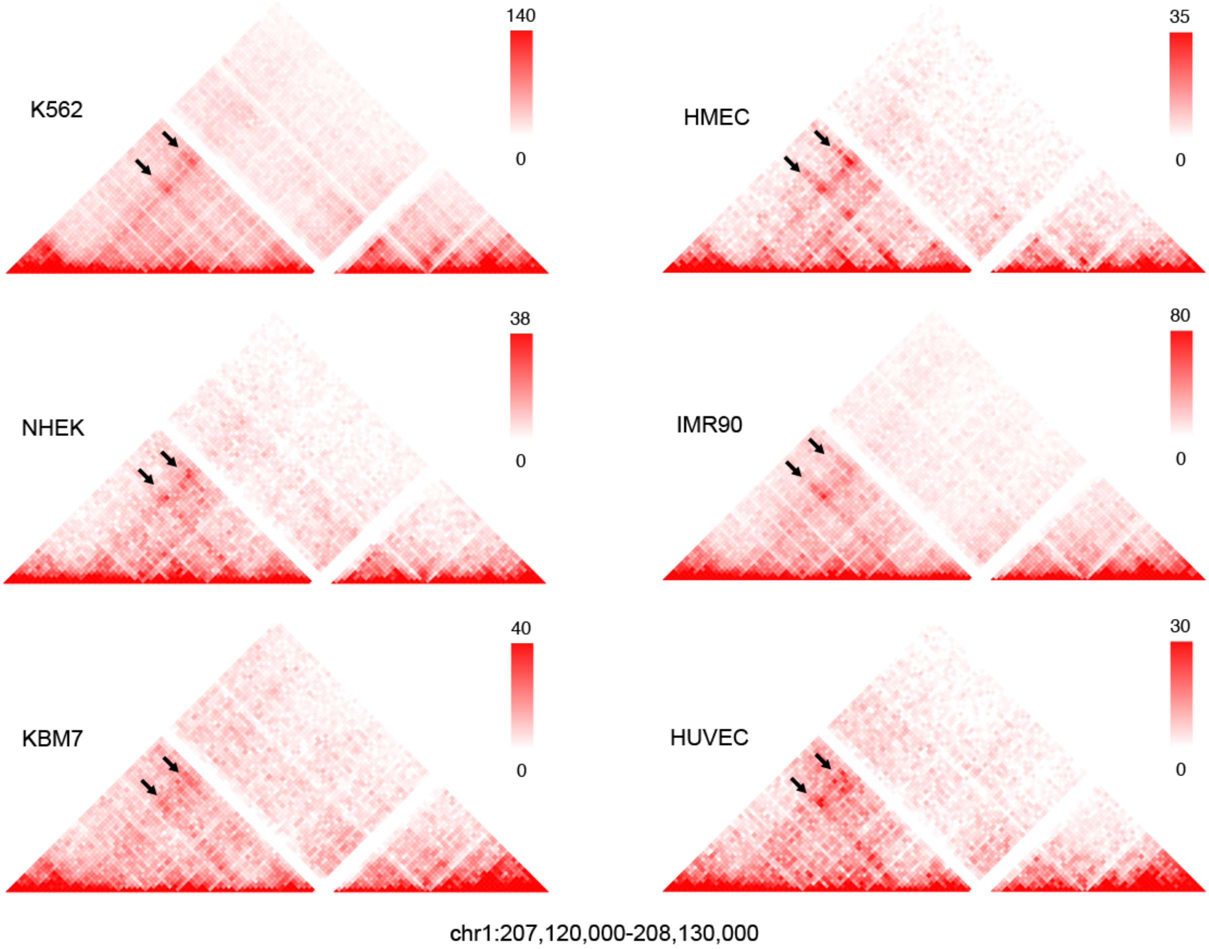


**Supplementary Figure 1: Pattern of interactions in Hi-C data across several cell lines were consistent and indicated that the RCA gene cluster may be divided into two TADs.**
Raw Hi-C data at 10 kb resolution from Rao *et al.* (1) for the 1 Mb region across the RCA genes were visualised using the 3D Genome Browser. Highly frequent chromatin interactions between upstream RCA genes (*C4BPB, C4BPA, CD55, CR2* and *CR1*) in GM12878 were observed in all cell lines assessed in this dataset (arrows), as identified in Figure 1.

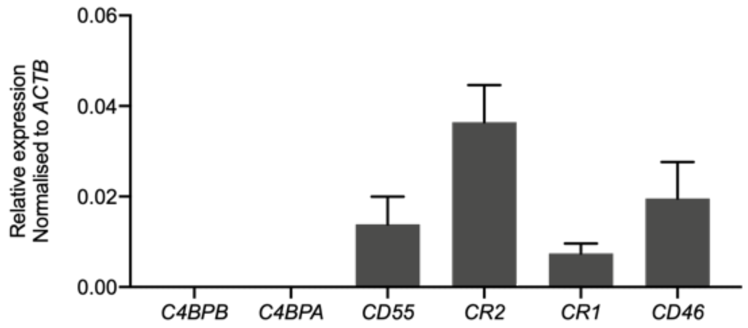


**Supplementary Figure 2: Receptor-bound RCA members are expressed in B cell line B-0028.**

Transcript abundance of RCA genes was measured by qPCR. Values were normalised to the β-actin gene (*ACTB*) using the ΔCt method. Bars represent mean relative expression ± SEM from 3 biological replicates.

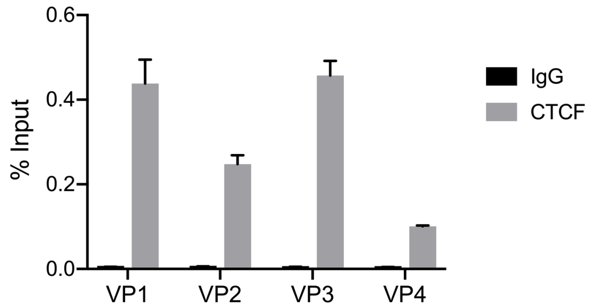


**Supplementary Figure 3: CTCF enrichment at 4C viewpoints in the B-0028 B lymphoblastoid cell line was confirmed by ChIP-qPCR using the percent input method.**
Grey bars indicate H3K27ac enrichment at the target locus, and black bars show enrichment using a non-specific IgG control antibody. All data are presented as mean ± SEM from at least 3 biological replicates.

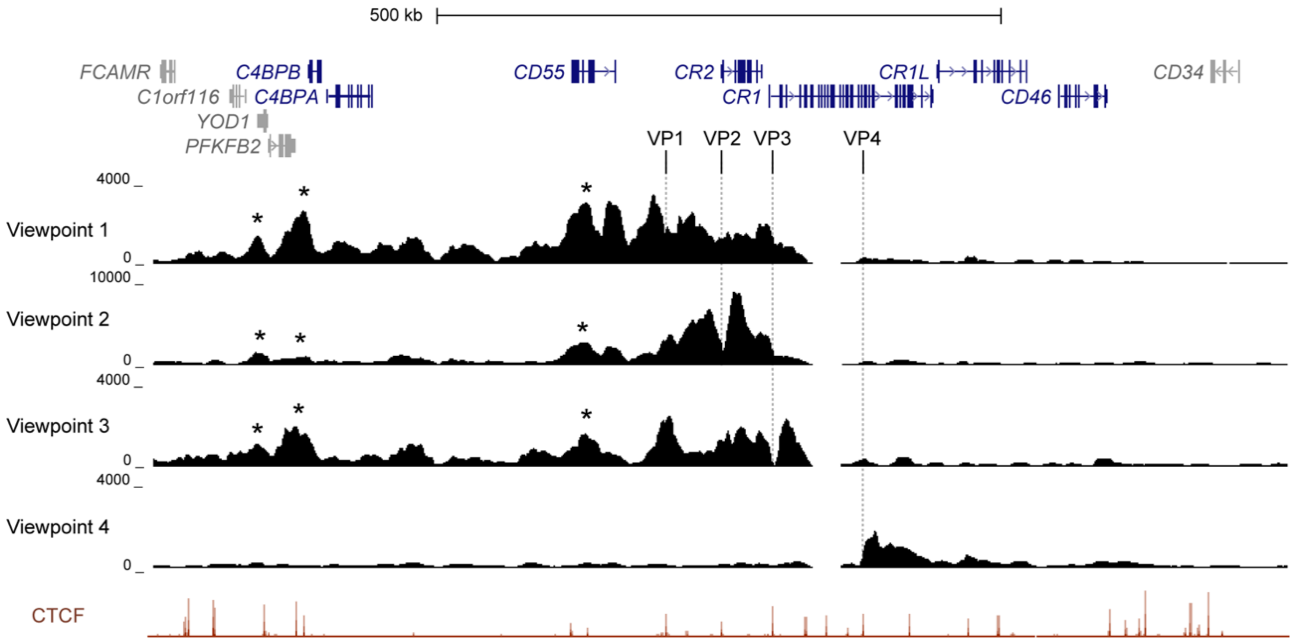


**Supplementary Figure 4: 4C-seq maps from CTCF viewpoints in the RCA gene cluster were replicated in the B-0056 lymphoblastoid cell line.**

Maps were generated from four viewpoints on CTCF sites, as indicated by GM12878 ChIP-seq signal for CTCF from ENCODE, in the intergenic region between *CR2* and *CD55* (viewpoint 1), intron 1 of *CR2* (viewpoint 2), intron 1 of *CR1* (viewpoint 3) and intron 29 of CR1 (viewpoint 4).

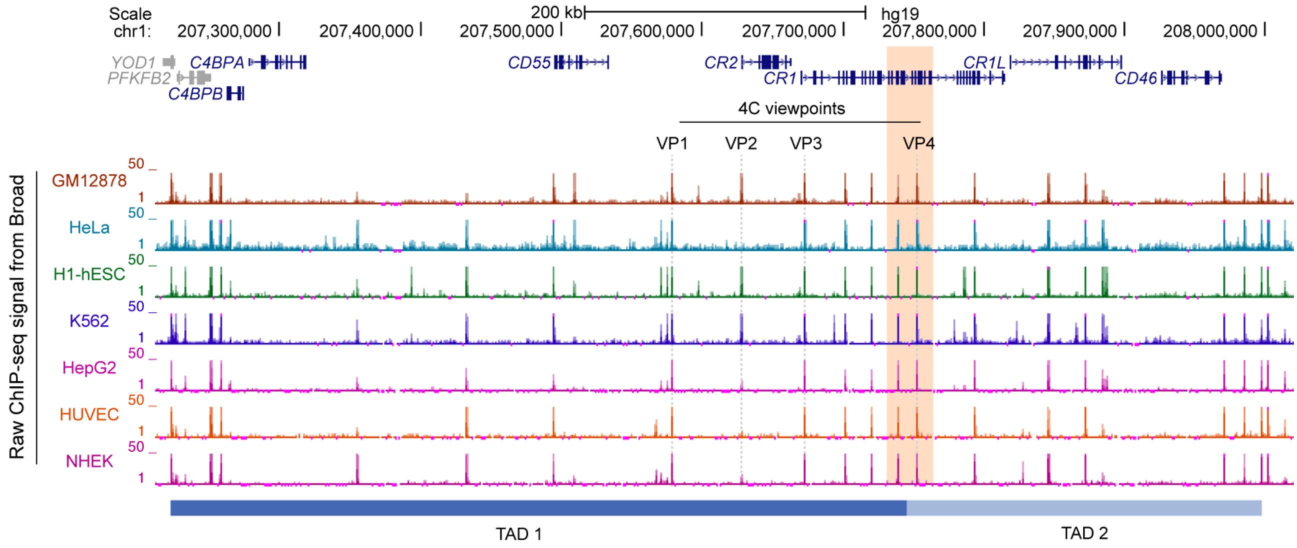


**Supplementary Figure 5: Divergent CTCF sites at the inter-TAD boundary (orange) were constitutive across multiple cell types.**

Raw ChIP-seq signal for CTCF from ENCODE/Broad was visualised using the UCSC Genome Browser on hg19. CTCF sites used as 4C viewpoints in TAD 1 (VP1, VP2, VP3) and TAD 2 (VP4) of the RCA gene cluster are indicated.

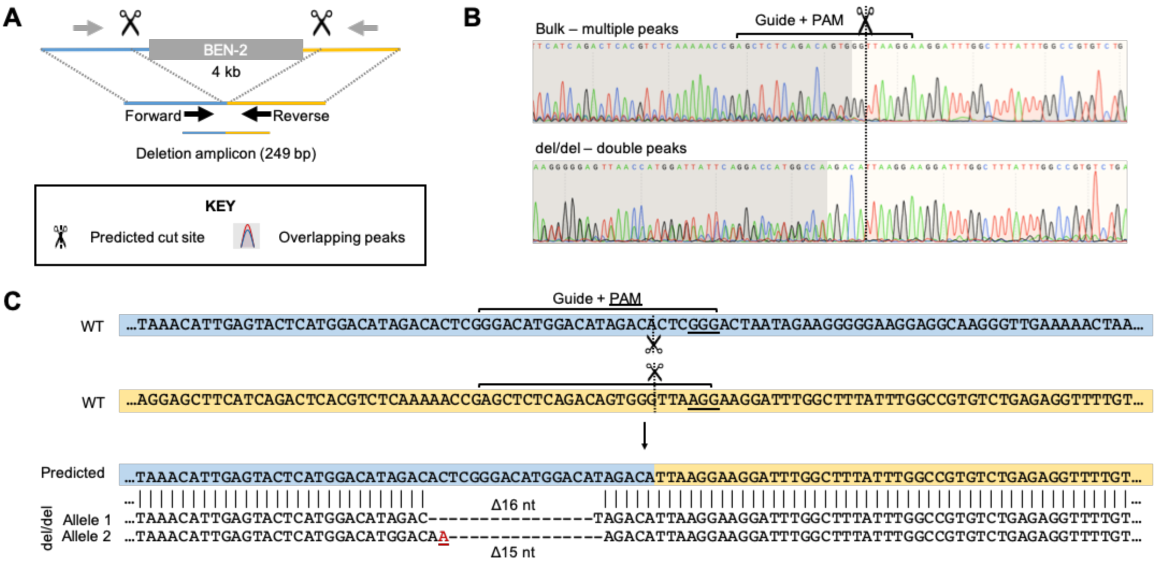


**Supplementary Figure 6**: **Sanger sequencing of deletion product amplified from BEN-2 CRISPR deletion clone confirms homozygous deletion genotype.**

A. Schematic of strategy using two Cas9 guides to delete the BEN-2 region and primer design to conduct initial screen of CRISPR clones.

B. Sanger sequencing chromatograms of the deletion product sequenced using the reverse primer (reverse-complement chromatograms are shown). Sequencing of the deletion product from the polyclonal bulk population revealed a region in close proximity to the predicted cut site containing multiple peaks, indicating that numerous alleles are present in the population. In contrast, the deletion product amplified from the monoclonal del/del population resulted in only single or double peaks, indicating that only two different alleles are present.

C. Sequence comparison of the predicted deletion amplicon and del/del clone confirmed the identity of the deletion product and successful homozygous deletion of the BEN-2 region. Poly Peak Parser (2) was used to call the sequence of double peaks in the Sanger chromatogram and sequence was manually examined to determine the resulting genotype. Indels which differed by one nucleotide were identified, which resulted in the double peaks after the Sanger sequencing.

### SUPPLEMENTARY TABLES

**Supplementary Table 1:** List of primers used.

| Primer | Sequence (5’ to 3’) |
|---|---|
| 4C VP1 read primer | TGCTTTTATGAAGGATTCTCTGATC |
| 4C VP2 read primer | CTACGGATGGTTAATGTTTAGGATC |
| 4C VP3 read primer | TTGGTTTCATCGAGTTGTGATC |
| 4C VP4 read primer | GTGTGACCCAGAATTCAGATC |
| 4C VP1 non-read primer | GTTGGTTGTGTGTATAAGGCG |
| 4C VP2 non-read primer | TCTTGGCTACAGGATGGGTC |
| 4C VP3 non-read primer | GAGGCATATAATTTATGTTCGTGT |
| 4C VP4 non-read primer | ACCATGCAGGATTTCTGGAG |
| 4C read adapter | AATGATACGGCGACCACCGAACACTCTTTCCCTACACGACGCTCTTC CGATCT |
| 4C non-read adapter | CAAGCAGAAGACGGCATACGA |
| BEN-1 forward FWD | CGGGGTACCAGTGCGCATGGGCTATTTACC |
| BEN-1 forward REV | GCGACGCGTTGAACCAGACCCAGGACTCAG |
| BEN-1 reverse FWD | GCGACGCGTAGTGCGCATGGGCTATTTACC |
| BEN-1 reverse REV | CGGGGTACCTGAACCAGACCCAGGACTCAG |
| BEN-2 forward FWD | CGGGGTACCTGAAGCCATCTCATCCCACAC |
| BEN-2 forward REV | GCGACGCGTACAGGCATGTGCCAAGTACAC |
| BEN-2 reverse FWD | GCGACGCGTTGAAGCCATCTCATCCCACAC |
| BEN-2 reverse REV | CGGGGTACCACAGGCATGTGCCAAGTACAC |
| BEN-3 forward FWD | GCGACGCGTTCCACAGAGCCAACAGCATTC |
| BEN-3 forward REV | GGAAGATCTAGCCAGAAGCACAGCTGTATG |
| BEN-3 reverse FWD | GGAAGATCTTCCACAGAGCCAACAGCATTC |
| BEN-3 reverse REV | GCGACGCGTAGCCAGAAGCACAGCTGTATG |
| BEN-4 forward FWD | CGGGGTACCTGCAGATGGAGGTTCCTAGAG |
| BEN-4 forward REV | GCGACGCGTTGTGCTGTTCATAAGCCATCC |
| BEN-4 reverse FWD | GCGACGCGTTGCAGATGGAGGTTCCTAGAG |
| BEN-4 reverse REV | CGGGGTACCTGTGCTGTTCATAAGCCATCC |
| *ACTB* mRNA FWD | ACCTTCTACAATGAGCTGCG |
| *ACTB* mRNA REV | CCTGGATAGCAACGTACATGG |
| *CD55* mRNA FWD | TGCAACCATCTCCTTCTCATG |
| *CD55* mRNA REV | GGTGCTGGACAATAAATTTCTCTG |
| *CR2* mRNA FWD | TGCCTGTAAAACCAACTTCTC |
| *CR2* mRNA REV | AGCAAGTAACCAGATTCACAG |
| *CD46* mRNA FWD | TCAGTAGCAATTTGGAGCGG |
| *CD46* mRNA REV | AGGTGCAGGATCACAACTATAAG |
| BEN-2 ChART FWD | AAAGTCCCATGCAACACTGG |
| BEN-2 ChART REV | AGCAAGGTTCAGAGATGTGC |
| 4C VP1 CTCF ChIP FWD | AGGCCATTGTCACACTGAAAC |
| 4C VP1 CTCF ChIP REV | GTGGTGACCCTGATGATGTG |
| 4C VP2 CTCF ChIP FWD | TAGCTTTGAGGGACCACTGC |
| 4C VP2 CTCF ChIP REV | AATTCTGGAGGTCCCAGCTC |
| 4C VP3 CTCF ChIP FWD | TTCATCCACAACAGCAGAGC |
| 4C VP3 CTCF ChIP REV | TGCCTGGTAAAGCTTAATTCG |
| 4C VP4 CTCF ChIP FWD | ACCACTGAGCTGGGAAGATG |
| 4C VP4 CTCF ChIP REV | TTTTGGTCAGCAGGATTGTG |
| BEN-2 H3K27ac ChIP FWD | GCAGCACATAAGGGTTCCAG |
| BEN-2 H3K27ac ChIP REV | GCAGGGCAGAAGAAGGAATG |
| CRISPR sub-cloning FWD | TGCTCTAGAGCTGGCCTTTTGCTCACATG |
| CRISPR sub-cloning REV | TTGGGTACCGCCATTTGTCTGCAGAATTG |
| CRISPR deletion FWD | TGCAGCTGGAAGCCATTATAC |
| CRISPR deletion REV | ATAGGGTCTCATTCTGTTGCC |
| CRISPR non-deletion FWD | AGGTTCAGGGAGTCTGAGGTG |
| CRISPR non-deletion REV | TTCTGCCCTTGTCCACCAAAC |

**Supplementary Table 2:** Hi-C sequencing data from Rao *et al.* used in mHi-C analysis.

| Cell line | Replicate | RE | Sequencing Depth | Dataset no. |
|---|---|---|---|---|
| GM12878 | rep2 | MboI | 314,309,612 | HIC019 |
| GM12878 | rep3 | MboI | 389,241,982 | HIC020 |
| GM12878 | rep4 | MboI | 178,301,500 | HIC021 |
| GM12878 | rep5 | MboI | 670,595,480 | HIC022 + HIC023 |
| GM12878 | rep6 | MboI | 111,656,957 | HIC024 |
| GM12878 | rep7 | MboI | 704,913,498 | HIC025 + HIC026 |
| GM12878 | rep8 | MboI | 328,197,634 | HIC027 |
| GM12878 | rep9 | MboI | 240,113,395 | HIC028 + HIC029 |
| GM12878 | rep32 | DpnII | 228,884,884 | HIC040 + HIC041 |
| GM12878 | rep33 | DpnII | 338,665,761 | HIC042 |

**Supplementary Table 3:** CRISPR gRNA sequences for enhancer deletion.

| Guide | Sequence (5’ to 3’) | Genomic co-ordinates (hg38) |
|---|---|---|
| g1 | GGACATGGACATAGACACTC | chr1:207411153-207411175 |
| g2 | AGCTCTCAGACAGTGGGTTA | chr1:207415757-207415779 |

**Supplementary Table 4:** Double elite putative enhancers in TAD 1 and TAD 2 of the RCA gene cluster, and their identified gene targets from GeneHancer.

| | GeneHancer ID | Relative location | Predicted gene targets |
|---|---|---|---|
| TAD 1 | GH01J207064 | *PFKFB2* | *C1orf116 FCAMR PIGR IL20 PFKFB2 YOD1 C4BPB* |
| | GH01J207066 | *PFKFB2* | *C1orf116 FCAMR PIGR IL20 PFKFB2 YOD1 C4BPB* |
| | GH01J207070 | *PFKFB2* | *PFKFB2 C1orf116 FCAMR PIGR C4BPA IL20 YOD1 C4BPB* |
| | GH01J207077 | *PFKFB2 – C4BPB* | *PFKFB2 CR1 C4BPA FCAMR PIGR C4BPB YOD1* |
| | GH01J207204 | *C4BPA – CD55* | *CD55 C4BPAP3 C4BPAP2* |
| | GH01J207239 | *C4BPA – CD55* | *CD55 CR2 IL19 C4BPA ENSG00000237074* |
| | GH01J207280 | *C4BPA – CD55* | *CD55 CR2 C4BPA ENSG00000237074* |
| | GH01J207298 | *C4BPA – CD55* | *CD55 ENSG00000237074* |
| | GH01J207304 | *C4BPA – CD55* | *CD55 CR2 ENSG00000237074* |
| | GH01J207306 | *C4BPA – CD55* | *CR2 CD55 ENSG00000237074* |
| | GH01J207309 | *C4BPA – CD55* | *CD55 EIF2D CR2 ENSG00000237074* |
| | GH01J207316 | *C4BPA – CD55* | *CDCA4P4 CD55 CD46 EIF2D ENSG00000234981 CR2 CR1 ENSG00000237074* |
| | GH01J207327 | *CD55* | *CD55 CR2 LOC105372881* |
| | *GH01J207333** | *CD55* | *CD55 EIF2D CR2 CR1 CD46 LOC105372881* |
| | GH01J207363 | *CD55 – CR2* | *EIF2D CD55 CR2 LOC105372881* |
| | GH01J207407 | *CD55 – CR2* | *CR1 CD55 CR2 C4BPB C4BPA CD46 YOD1 ENSG00000283044 LOC105372881* |
| | *GH01J207411** | *CD55 – CR2* | *CR2 CR1 CD55 ENSG00000283044 LOC105372881* |
| | *GH01J207424** | *CD55 – CR2* | *CR2 CR1 C1orf116 C4BPA LOC105372880* |
| TAD 2 | GH01J207625 | *CR1* | *CR1 C4BPA CD46P1 ENSG00000236911 GC01M207468* |
| | GH01J207648 | *CR1L* | *CR1L C4BPA MIR29B2CHG CD46P1 CDCA4P3 GC01M207468* |
| | GH01J207649 | *CR1L* | *CR1L C4BPA MIR29B2CHG CR2 CD46P1 CDCA4P3 GC01M207468* |
| | GH01J207755 | *CD46* | *CD46 MIR29B2CHG CDCA4P4* |

* These predicted enhancers were functionally investigated.

**Supplementary Table 5:** Strong candidate B cell enhancers (BENs) were predicted to regulate genes on GeneHnacer using eQLT analyses, promoter interactions from capture Hi-C (CHi-C) and/or enhancer RNA and mRNA co-expression (eRNA).

| | | | BEN-1 | BEN-2 | BEN-3 | BEN-4 |
|---|---|---|---|---|---|---|
| Non-RCA | *PIGR* | Method | | | | CHi-C |
| | | Score | | | | 9.7 |
| | *FCAMR* | Method | | | | CHi-C |
| | | Score | | | | 9.7 |
| | *C1orf116* | Method | | | CHi-C | |
| | | Score | | | 10.2 | |
| | *PFKFB2* | Method | | | | |
| | | Score | | | | |
| RCA | *C4BPA* | Method | | | CHi-C | |
| | | Score | | | 9.4 | |
| | *CD55* | Method | eQTL<br>eRNA | CHi-C | | CHi-C |
| | | Score | *27.2** | 10.3 | | 10.4 |
| | *CR2* | Method | eQTL | CHi-C<br>eRNA | CHi-C<br>eRNA | CHi-C |
| | | Score | 9.6 | *18.7** | *18.1** | 12.3 |
| | *CR1* | Method | eRNA | CHi-C | CHi-C | CHi-C |
| | | Score | 7.2 | 11.8 | 11.1 | 11.1 |
| | *CR1L* | Method | | | | |
| | | Score | | | | |
| | *CD46* | Method | eRNA | | | |
| | | Score | 3.6 | | | |

* These predicted gene-enhancer interactions were identified using more than one method on GeneHancer.
